# Same role but different actors: genetic regulation of post-translational modification of two distinct proteins

**DOI:** 10.1101/2021.05.04.442584

**Authors:** Arianna Landini, Irena Trbojević-Akmačić, Pau Navarro, Yakov A. Tsepilov, Sodbo Z. Sharapov, Frano Vučković, Ozren Polašek, Caroline Hayward, Tea Petrovic, Marija Vilaj, Yurii S. Aulchenko, Gordan Lauc, James F. Wilson, Lucija Klarić

## Abstract

Post-translational modifications (PTMs) diversify protein functions and dynamically coordinate their signalling networks, influencing most aspects of cell physiology. Nevertheless, their genetic regulation or influence on complex traits is not fully understood. Here, we compare for the first time the genetic regulation of the same PTM of two proteins – glycosylation of transferrin and immunoglobulin G (IgG). By performing genome-wide association analysis of transferrin glycosylation, we identified 10 significantly associated loci, all novel. Comparing these with IgG glycosylation-associated genes, we note protein-specific associations with genes encoding glycosylation enzymes (transferrin - *MGAT5, ST3GAL4, B3GAT1*; IgG - *MGAT3, ST6GAL1*) as well as shared associations (*FUT6, FUT8*). Colocalisation analyses of the latter suggest that different causal variants in the FUT genes regulate fucosylation of the two proteins. We propose that they affect the binding of different transcription factors in different tissues, with fucosylation of IgG being regulated by IKZF1 in B-cells and of transferrin by HNF1A in liver.

## Introduction

Post-translational modifications (PTMs) are essential mechanisms used by cells to diversify and extend their protein functions beyond what is dictated by protein-coding sequences in the genome. These chemical reactions range from the addition of small moieties, such as phosphate (phosphorylation), complex biomolecules, as in glycosylation, to proteolytic cleavage^1^. PTMs alter the structure and properties of proteins and are thus involved in the dynamic regulation of most cellular events. It is common for a PTM enzyme to target multiple substrates or interact with multiple sites. For example, only 18 histone deacetylases target more than 3600 acetylation sites on 1750 proteins^2^. Environmental or pathological conditions can lead to dysregulation of PTM activities, which has been related to aging^3^ and several diseases, including cancer, diabetes, and neurodegeneration^4–10^. Despite their importance, little is known about genetic regulation of post-translational modifications.

N-glycosylation is one of the most common protein PTMs, where carbohydrate structures called glycans are covalently attached to an asparagine (Asn) residue of a polypeptide backbone. N-glycans are characterised by vast structural diversity and high complexity. While polypeptides are encoded by a single gene, N-glycan structures result from a sophisticated interplay of glycosyltransferases, glycosidases, transporters, transcription factors, and other proteins^11^. Protein N-glycosylation is involved in a multitude of biological processes^12^. Accordingly, changes in N-glycosylation patterns have been associated with aging^13^ and a wide range of diseases, including Parkinson’s disease^14^, lower back pain^15^, rheumatoid arthritis^16^, ulcerative colitis^17^, Crohn’s disease^17^, type 2 diabetes^18^ and cancer^19–21^. In addition, N-glycans are considered as potential therapeutic targets^22^ and prognostic biological markers^18,23–25^.

As with other PTMs, genetic regulation of N-glycosylation is not yet fully understood. Previous genome-wide association studies (GWAS) have so far focused either on the N-glycome of total blood plasma proteins as a whole or on glycosylation of one specific protein - immunoglobulin G (IgG)^26–33^. IgG antibodies are one of the most abundant proteins in human serum, and their alternative N-glycosylation is suggested to trigger different immune response and thus impacts the action of the immune system^34^. N-glycan structures are predominantly of the biantennary complex type and vary due to additions of core fucose, galactose, sialic acid, and bisecting N-acetylglucosamine (GlcNAc), with disialylated digalactosylated biantennary glycan with core fucose and bisecting GlcNAc being the most complex N-glycan structure on IgG^35^. While a clear overlap in genetic control between total plasma proteins and IgG N-glycosylation was highlighted by previous studies^28^, it was not possible, until now, to identify protein-specific N-glycosylation pathways for glycoproteins other than IgG due to technical challenges hampering isolation of other glycoproteins in large cohorts.

Here we report genes associated with the regulation of transferrin N-glycosylation and compare these with the genetic regulation of glycosylation of a different protein (IgG). Transferrins are blood plasma glycoproteins regulating the level of iron in an organism. Iron plays a central role in many essential biochemical processes of human physiology: the cells’ need for iron in the face of potential danger as an oxidant has given rise to a complex system that tightly regulates iron levels, tissue distribution, and bioavailability^36^. Human transferrin has two N-glycosylation sites – at the N432 and N630 residues, with biantennary disialylated digalactosylated glycan structure without fucose being the most abundant glycan attached^37,38^. We performed, for the first time, genome-wide association meta-analysis (GWAMA) of 35 transferrin N-glycan traits (N=1890) and compared it with GWAMA of 24 IgG N-glycan traits (N=2020) in European-descent cohorts, discovering both protein-specific and shared associations. For loci associated with the N-glycosylation PTM of both transferrin and IgG, we used colocalisation analysis to assess whether the underlying causal variants are protein-specific or rather shared between these proteins. We then suggested a molecular mechanism by which these independent causal variants could regulate the expression of glycosylation related genes in different tissues. To the best of our knowledge, this is the first study investigating whether the same PTM of two proteins is regulated by the same genes and whether they are driven by the same causal genetic variants.

## Results

### Loci associated with transferrin N-glycosylation

To investigate the genetic control of transferrin N-glycosylation and assess whether the same genes and underlying causal variants are associated with N-glycosylation of both transferrin and IgG, we first performed GWAS of glycosylation for each protein (i.e. transferrin and IgG). A more extensive GWAS on the genetic regulation of IgG glycosylation has already been published^30^, so we focus here on glycosylation of transferrin. We performed GWAS of 35 ultra-high-performance liquid chromatography (UHPLC)-measured transferrin N-glycan traits and Haplotype Reference Consortium (HRC) r1.1-imputed genetic data in two cohorts of European descent (N=1890). To identify secondary association signals at each genomic region, we performed approximate conditional analysis on transferrin N-glycan traits using GCTA-COJO software^39^. Overall, we identified 26 independently contributing variants, located in 10 genomic loci significantly associated (p-value ≤ 1.43×10^-9^, Bonferroni adjusted for the number of glycan traits) with at least one of the 35 transferrin N-glycan traits (Table 1, Figure 1, complete list of all associations in Supplementary Table 1). Multiple SNPs independently contributed to transferrin N-glycans variation in 6 out of 10 loci, all mapping to glycosyltransferase genes, plus the transferrin (*TF*) gene. The highest number of independently associated SNPs (7) was observed for the sialyltransferase locus, *ST3GAL4*, followed by 4 SNPs in the acetylglucosaminyltransferase locus, *MGAT5*, and in the glucuronyltransferase locus, *B3GAT1*. Lastly, 2 SNPs independently contributed to transferrin glycosylation in the fucosyltransferase loci, *FUT8* and *FUT6*, and also the transferrin (*TF*) locus itself (Supplementary Table 2).

**Table 1.**
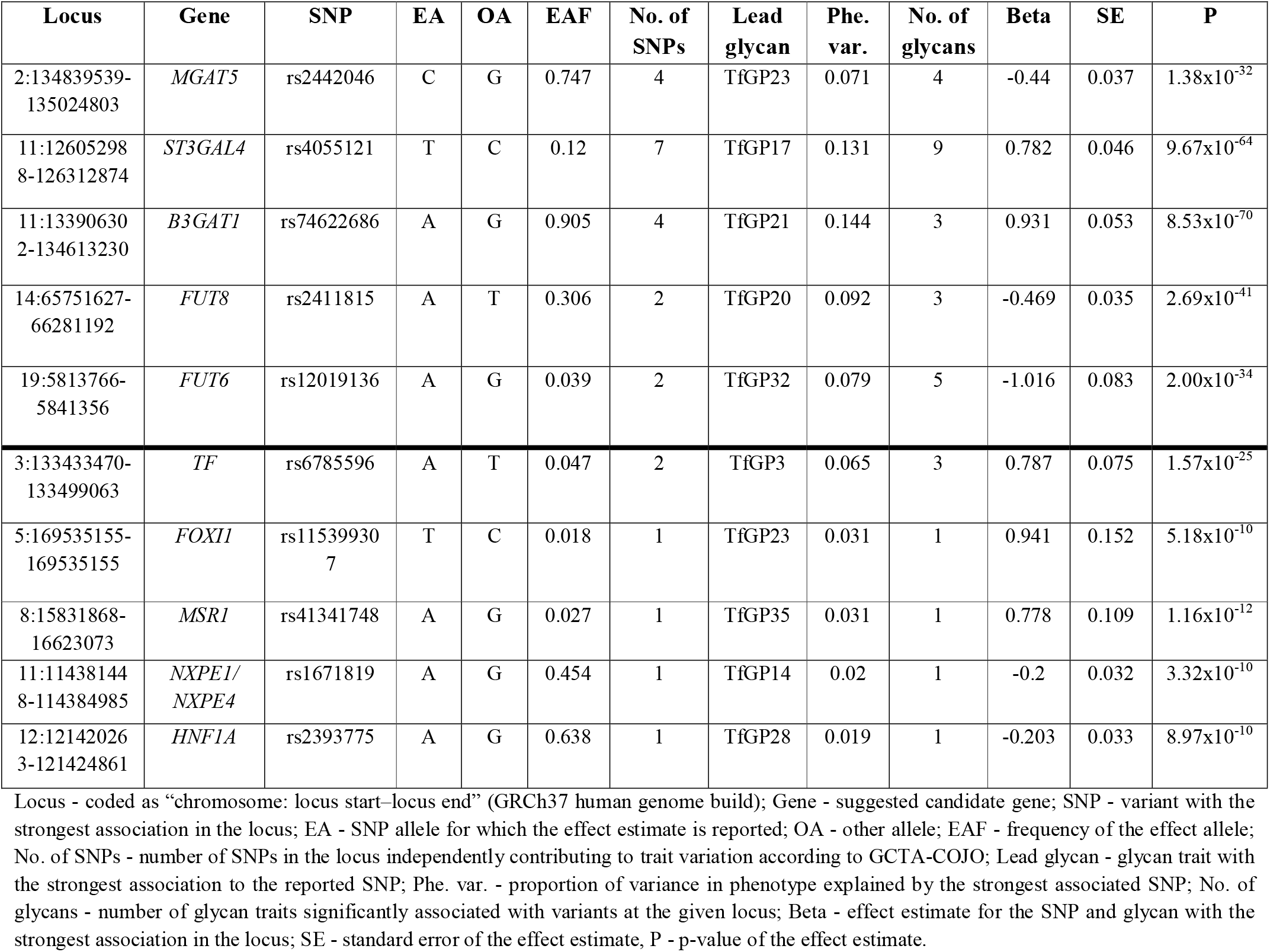
Loci genome-wide significantly associated with at least one of the 35 transferrin N-glycan traits in GWAMA. Glycosyltransferase loci are reported at the top of the table, while other loci are listed at the bottom of the table. Each locus is represented by the SNP with the strongest association in the region.

**Figure 1.**
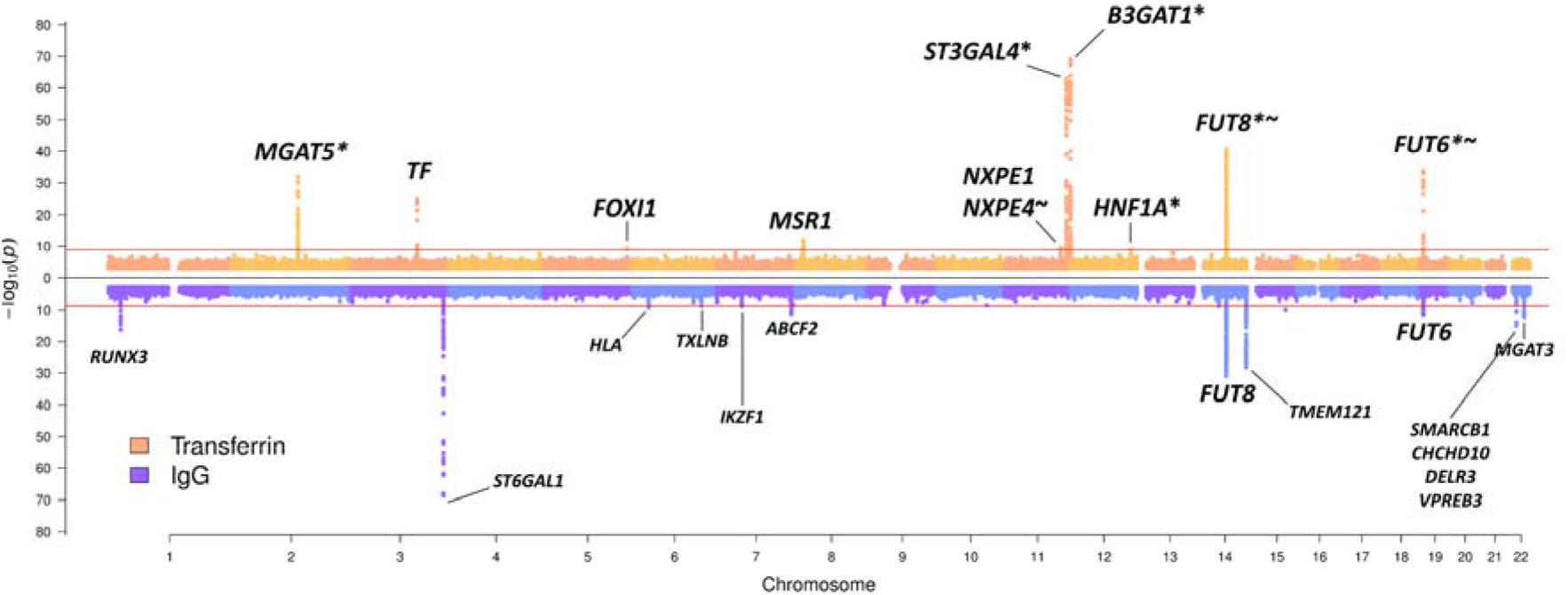
Transferrin and IgG N-glycome GWAMA summary Miami plot. Miami plot pooling together meta-analysis results obtained across all 35 transferrin glycan traits at the top in orange, and across all 24 IgG glycan traits at the bottom in blue. For transferrin N-glycome associations, * marks loci previously reported in total blood plasma N-glycome GWAS^26–28^, while ~ marks loci previously reported in IgG N-glycome GWAS^30–32,40^. For simplicity, SNPs with p-value > 1×10^-3^ are not plotted. The Bonferroni-corrected genomewide significance threshold for the transferrin N-glycome meta-analysis (horizontal red line in the top part of the plot) corresponds to 1.43×10^-9^, while the Bonferroni-corrected genomewide significance threshold for the IgG N-glycome meta-analysis (horizontal red line in the bottom part of the plot) corresponds to 2.08×10^-9^. Gene or sets of genes annotated for transferrin N-glycome loci have been prioritised in this study; gene or sets of genes annotated for IgG N-glycome loci are those prioritised by Klarić et al.^30^

### Prioritising candidate genes associated with transferrin N-glycosylation

For the 10 loci associated with the transferrin N-glycome, we identified plausible candidate genes following multiple lines of evidence, such as evaluating the biological role of the candidate gene in the context of protein N-glycosylation, assessing SNP pleiotropy with eQTLs, and investigating variant effects on the coding sequence or on putative transcription factor binding sites.

#### Positional mapping and biological role

The majority of genes that were closest to transferrin N-glycosylation-associated variants had a clear biological link to protein N-glycosylation. In particular, for 5 out of 10 loci the closest genes (i.e. *MGAT5, ST3GAL4, B3GAT1, FUT8*, and *FUT6*) encode glycosyltransferases, key enzymes in protein glycosylation, that have been previously associated with IgG and/or total plasma protein glycosylation (Supplementary Table 3). Another gene closest to transferrin N-glycosylation-associated variants and with a validated functional role in plasma proteins glycosylation is *HNF1A*, a transcription factor previously associated with protein fucosylation (Supplementary Table 3). On the other hand, we also identified 3 loci that had not been associated with N-glycosylation. A locus on chromosome 3 contains the transferrin (*TF*) gene, which encodes the transferrin glycoprotein. A locus on chromosome 5 contains *FOXI1*, encoding a member of the forkhead family of transcription factors (Forkhead box I1). Finally, a locus on chromosome 8 contains the *MSR1* gene, encoding the class A macrophage scavenger receptor, a trimeric integral membrane glycoprotein. Another gene of potential biological relevance at the chromosome 8 locus is the tumour suppressor candidate 3 (*TUSC3* which encodes a protein localised to the endoplasmic reticulum and acting as a component of the oligosaccharyltransferase complex, responsible for N-linked protein glycosylation.

#### Overlap and colocalisation with eQTL

Using eQTL analysis in PhenoScanner, transferrin N-glycan-associated genetic variants (and their proxies, LD r^2^ > 0.8) were identified to be significantly associated with the expression of multiple genes in several human tissues involved in transferrin metabolism (Supplementary Table 4a). For example, transferrin glycosylation variants were associated with *ST3GAL4* expression in liver and whole blood, with *B3GAT1* expression in visceral adipose omentum, liver, and whole blood, with *TF* expression in several adipose tissues and with *HNF1A, FUT8*, and *MGAT5* expression in whole blood. The majority of these genes were also the closest to the strongest association in the locus. We next used Summary data-based Mendelian Randomization (SMR) analysis followed by the Heterogeneity in Dependent Instruments (HEIDI) test^41^ to assess whether expression of these genes colocalises with transferrin glycosylation (TfGP) traits. SMR-HEIDI provided evidence of pleiotropy, suggesting that the same underlying causal SNPs are likely to regulate both transferrin glycosylation traits and gene expression, for *B3GAT1* in liver and peripheral blood and *ST3GAL4* in liver (Supplementary Table 4b).

#### Analysis of possible effects on amino acid sequence

We next explored whether any of the SNPs independently contributing to transferrin glycosylation (or their proxies) result in a change of amino acid sequence using the Ensembl Variant Effect Predictor (VEP)^42^. While the majority of associated variants (> 60%) were classified as intronic, several SNPs were identified as missense variants: rs115399307 (5_169535155_T/C) causes the substitution of the non-polar, aliphatic amino acid isoleucine (I) to the polar, hydrophilic amino acid threonine (T) in the FOXI1 transcription factor. Similarly, *NXPE4* variant rs550897 (11_114442103_A/G, r^2^=0.94 with rs1671819) causes an amino acid substitution from tyrosine (Y) to histidine (H), while *FUT6* variant rs17855739 (19_5831840_T/C, r^2^=0.95 with rs12019136) encodes a change from negatively charged glutamic acid (E) to positively charged lysine (K), which leads to a full-length, but inactive, enzyme^43^. Genetic variant rs41341748 (8_16012594_A/G) disrupts a stop codon sequence in *MSR1*, causing an elongated transcript with the amino acid arginine (Arg) added to the protein chain (Supplementary Table 5).

#### Analysis of possible effects on transcription factor binding sites

Finally, we used the regulatory sequence analysis tools (RSAT)^44^ to assess if transferrin N-glycosylation-associated genetic variants overlap transcription factor-binding sites (TFBSs) and are likely to affect transcription factor (TF) binding. From the list of prioritised genes, we selected the two encoding transcription factors, FOXI1 and HNF1A, and checked whether associated variants in the remaining 8 loci were likely to affect their binding. Overall, binding of both FOXI1 and HNF1A transcription factors is likely to be affected by the sentinel variant (the SNP with lowest p-value in the region for the given glycan trait) in the *FUT8* gene. In addition, binding of HNF1A is likely to be affected also by the sentinel variants in the *TF* and *ST3GAL4* loci (Supplementary Table 6).

### Shared genetic associations with complex traits and diseases

To assess whether transferrin glycosylation variants were also associated with complex traits and diseases we used PhenoScanner^45^, followed by SMR-HEIDI to determine whether the shared associations are caused by the same underlying causal variant (pleiotropy). We observed an overlap of transferrin N-glycan-associated SNPs and their proxies with variants associated with complex trait- and disease-associated variants for 5 out of 10 glycosylation loci (Supplementary Table 7a). Glycosylation SNPs at the *NXPE1/NXPE4* locus were pleiotropic with ulcerative colitis, and those from the *HNF1A* locus with C-reactive protein levels, LDL and total cholesterol (Supplementary Table 7b). For the remaining shared associations, we had no power to assess pleiotropy (Supplementary Results for further details). Interestingly, variants at the *TF* locus have been previously associated with serum concentration of carbohydrate-deficient transferrins (CDT) (Supplementary Table 7a), less glycosylated transferrin isoforms traditionally used as a biomarker of excessive alcohol consumption^46^, thus corroborating our finding for a related trait.

### Comparison of genetic regulation of glycosylation of transferrin and immunoglobulin G

One of the main aims of this study is to understand if the N-glycosylation of two proteins is regulated by the same enzymes and if so, whether the same underlying genetic variant or a set of variants are driving the process. To address this question, in addition to the already described GWAMA of transferrin glycosylation, we performed a GWAMA of 24 UHPLC IgG N-glycan traits in the same individuals (N=2020), following the same protocol. 13 loci were significantly associated with at least one of the 24 IgG N-glycan traits (Figure 1, Supplementary Table 8). The IgG N-glycome GWAS was annotated using genes or sets of genes prioritised by Klarić et al.^30^ By comparing the two GWAS we discovered mainly protein-specific associations, but also two genomic regions that were associated with glycosylation of both proteins (Figure 1). The protein-specific associations were with genes encoding known glycosylation enzymes (transferrin - *MGAT5, ST3GAL4, B3GAT1*; IgG - *ST6GAL1, MGAT3*), but also with transcription factors (transferrin - *HNF1A, FOXI1*; IgG - *IKZF1, RUNX3*), the protein itself (transferrin - *TF*; IgG - *TMEM121*, gene in proximity of *IGH* genes encoding immunoglobulin heavy chains) as well as other genes (transferrin - *MSR1*; IgG - *TXLNB, ABCF2, SMARCB1* region, HLA-region). Interestingly, the regions containing *FUT8* and *FUT6*, genes encoding fucosyltransferases, enzymes adding core and antennary fucose, respectively, to the synthetized glycan, were associated with glycosylation of both proteins (Figure 1). We then proceeded to assess whether the same underlying causal variants in these regions are controlling the process for both proteins using colocalisation analysis.

Given that multiple glycan traits of the same protein can be associated with the same locus, we first asked whether all glycan traits of the same protein associated with a certain locus, colocalise (Supplementary Figure 1). Indeed, we found strong support for colocalisation (PP.H4 > 80 %, where PP.H4 represents the posterior probability for the same underlying causal variant contributing to trait variation), suggesting that for a given protein, all glycan traits associated with these loci are regulated by the same underlying causal variant (Supplementary Table 9, Supplementary Figure 2-4). One example of within-protein colocalisation can be seen in Figure 2. We next tested whether at the same genomic region, glycosylation of two different proteins is regulated by the same underlying causal variants. For this, we selected as the protein-representative glycan trait the one with the lowest p-value in the given region (one pair for each locus - transferrin TfGP20 and IgG GP7 for the *FUT8* locus and transferrin TfGP32 and IgG GP20 for the *FUT6* locus) and proceeded to test for colocalisation between glycosylation of the two proteins. We found strong support against colocalisation in both genomic regions (PP.H3 = 100% at *FUT8* locus, PP.H3 = 99.71% at *FUT6* locus, where PP.H3 represents the posterior probability for different underlying causal variants contributing to trait variation) (Figure 3 and Figure 4, Supplementary Table 10). Since colocalisation methods are sensitive to multiple independent variants in the region contributing to the trait variation, which was the case here, we validated our findings with the PwCoCo approach^47^ (Methods) and again, obtained robust evidence against the colocalisation hypothesis for all tested traits in both loci (Supplementary Table 10 and Supplementary Results for further details).

**Figure 2.**
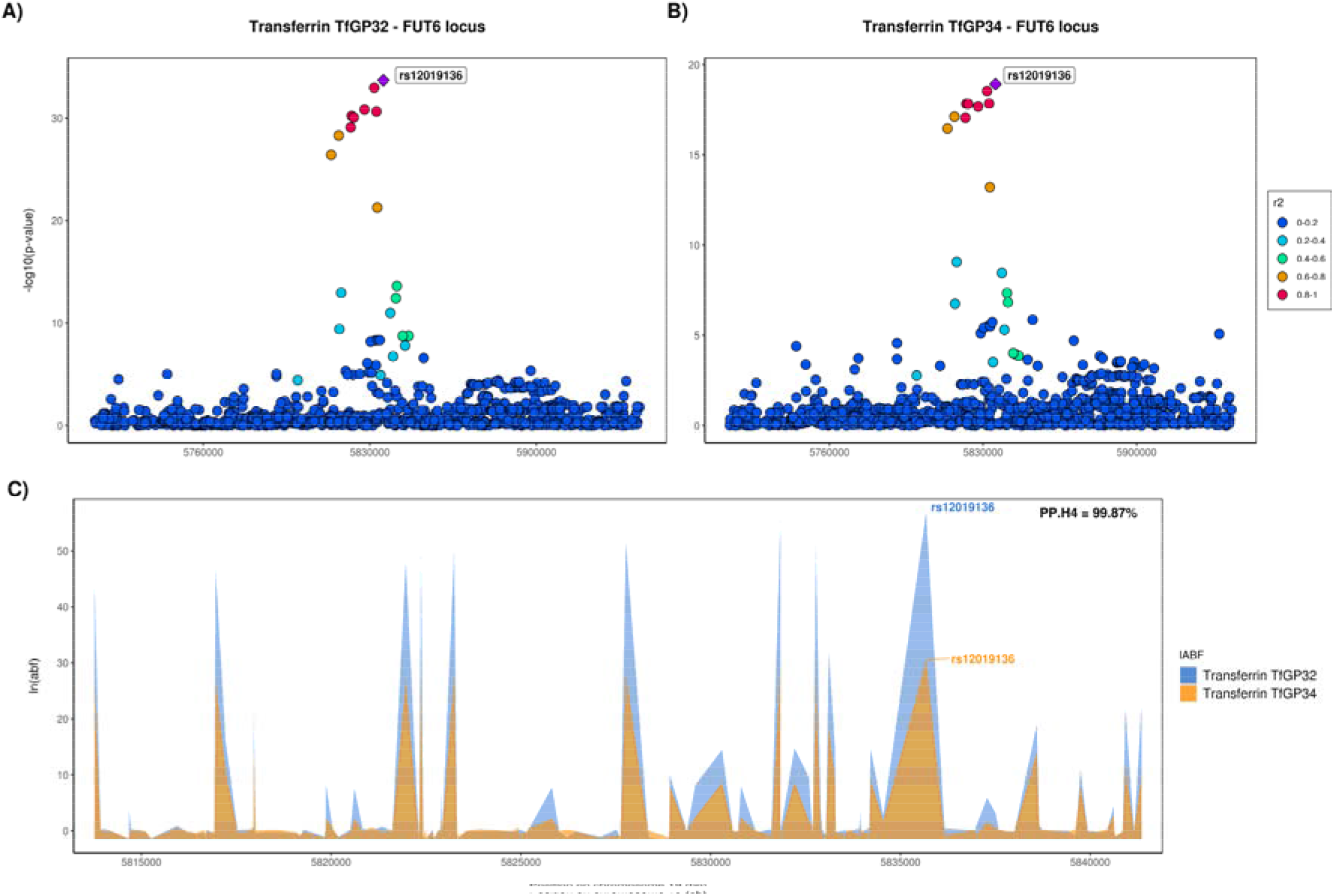
Local association patterns for transferrin (A) TfGP32 and (B) TfGP34 glycans, and (C) their colocalisation pattern at the *FUT6* locus. TfGP32 and TfGP34 association patterns colocalise, with PP.H4 (posterior probability for hypothesis 4, of colocalisation) of 99.87%. c) The logarithm of Approximate Bayes Factor (ABF) of each SNP for transferrin TfGP32 and transferrin TfGP34 in the *FUT6* region shows that TfGP32 and TfGP34 associations are concordant (the patterns of ln(ABF) calculated for each SNP of both traits overlap), suggesting that the same underlying causal variant is associated with both traits.

**Figure 3.**
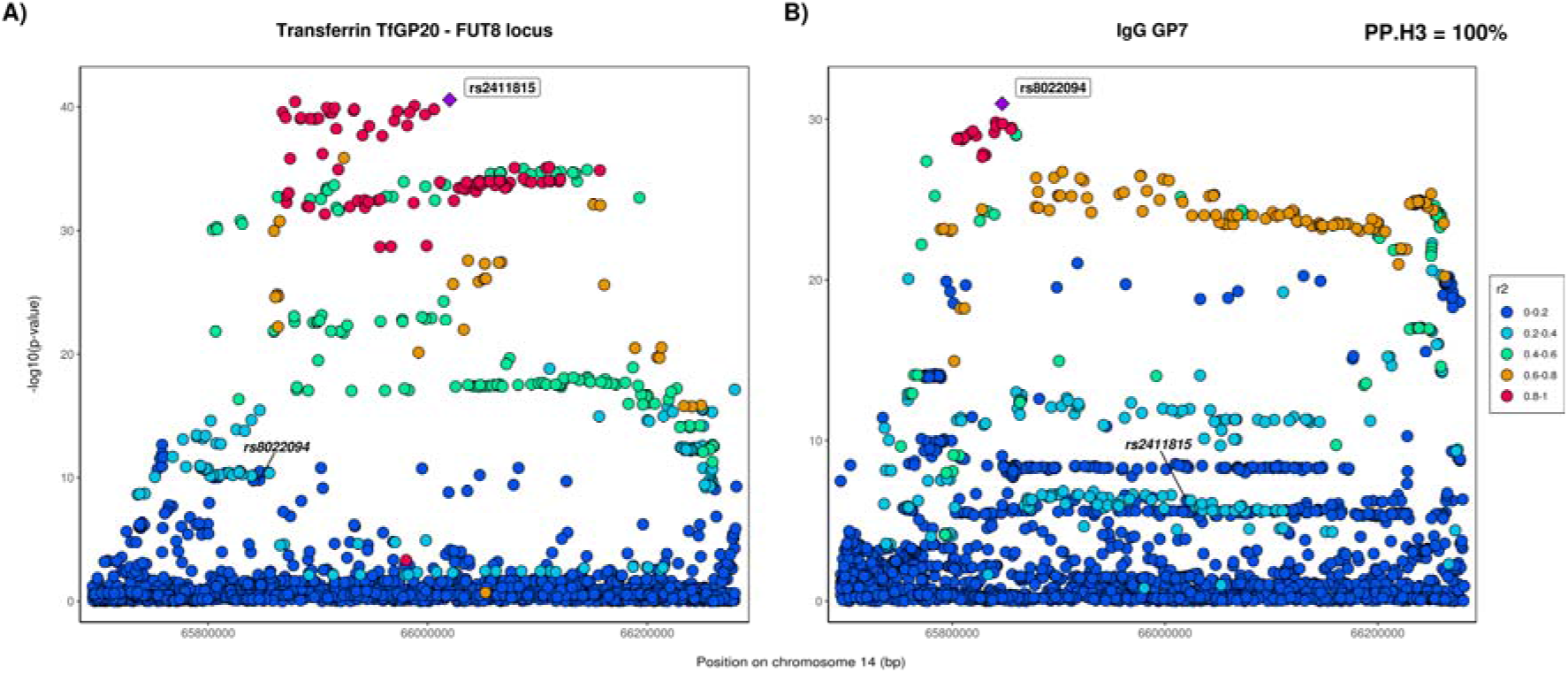
Local association patterns for (A) transferrin TfGP20 and (B) IgG GP7 glycans at the *FUT8* locus. TfGP20 and IgG GP7 association patterns do not colocalise (PP.H3 = 100% - posterior probability for hypothesis 3, of different causal variants). Colocalisation patterns are not reported since the width of the *FUT8* region makes the plot non-informative.

**Figure 4.**
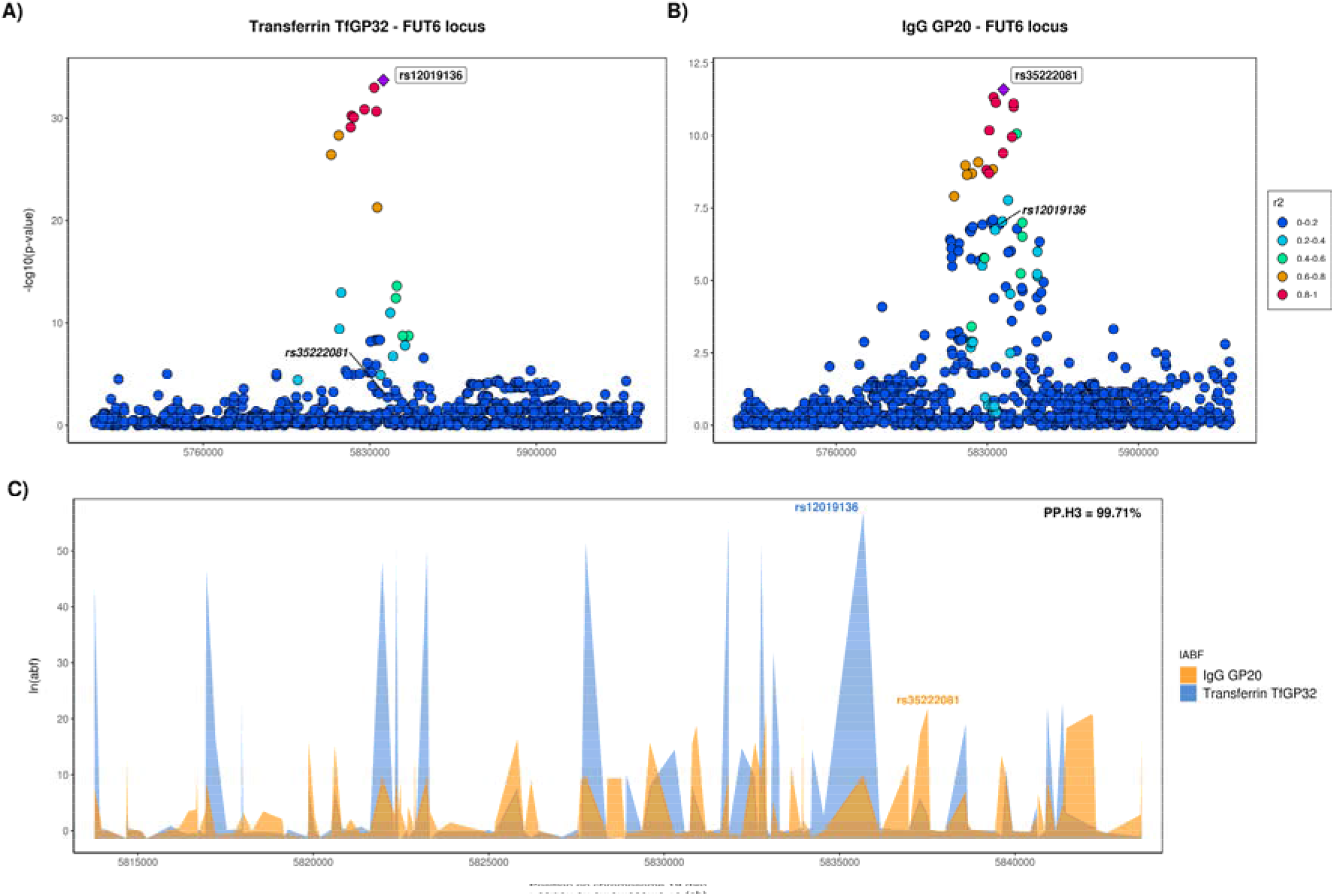
Local association patterns for (A) transferrin TfGP32 and (B) IgG GP20 glycans, and (C) their colocalisation pattern at the *FUT6* locus. TfGP32 and IgG GP20 association patterns do not colocalise (PP.H3, posterior probability for hypothesis 3, of different causal variants = 99.7%). The logarithm of Approximate Bayes Factor (ABF) of each SNP for transferrin TfGP32 and IgG GP20 in the *FUT6* region shows that TfGP32 and GP20 associations are not concordant (the patterns of ln(ABF) calculated for each SNP of both traits do not overlap), suggesting that two different underlying causal variants in this region regulate glycosylation of these two proteins.

Having established that different underlying causal variants regulate glycosylation at the *FUT6* and *FUT8* loci, we next explored the potential mechanisms behind these associations. The sentinel transferrin glycosylation SNP in the *FUT8* region is likely to affect binding of the HNF1A transcription factor (Supplementary Table 6) and it was previously shown that sentinel IgG glycosylation SNP in the same region potentially affects binding of the IKZF1 transcription factor^30^. In addition, we observed protein-specific associations with two transcription factors: transferrin glycosylation was associated with variants in the *HNF1A* locus and IgG glycosylation was associated with variants in the *IKZF1* locus (Figure 1). We therefore checked expression of these genes in tissues where the two proteins are predominantly expressed. It is known that plasma transferrin, encoded by *TF* gene, is mostly secreted by hepatocytes^48^, while IgG, the heavy chain constant region of which is encoded by *IGHG* gene, is predominantly synthesised by the antibody-secreting plasma cells, the fully differentiated form of B-lymphocytes^49^. Indeed, we see that *IGHG1* (encoding the most prevalent IgG1 subclass) is highly expressed in plasma cells and has low expression in hepatocytes, while the converse is true for *TF* (Figure 5). Similarly, the transcription factor encoded by *HNF1A* is predominantly expressed in the hepatocytes, while *IKZF1* is mainly expressed in plasma cells (Figure 5). Altogether these suggest that two distinct causal variants regulating glycosylation of transferrin and IgG in the *FUT8* locus are likely to have tissuespecific effects, where the transferrin-associated variant affects the binding of HNF1A in liver and the IgG-associated variant affects the binding of IKZF1 in plasma cells, with both influencing expression of the *FUT8* gene and therefore affecting fucosylation of the two proteins.

**Figure 5.**
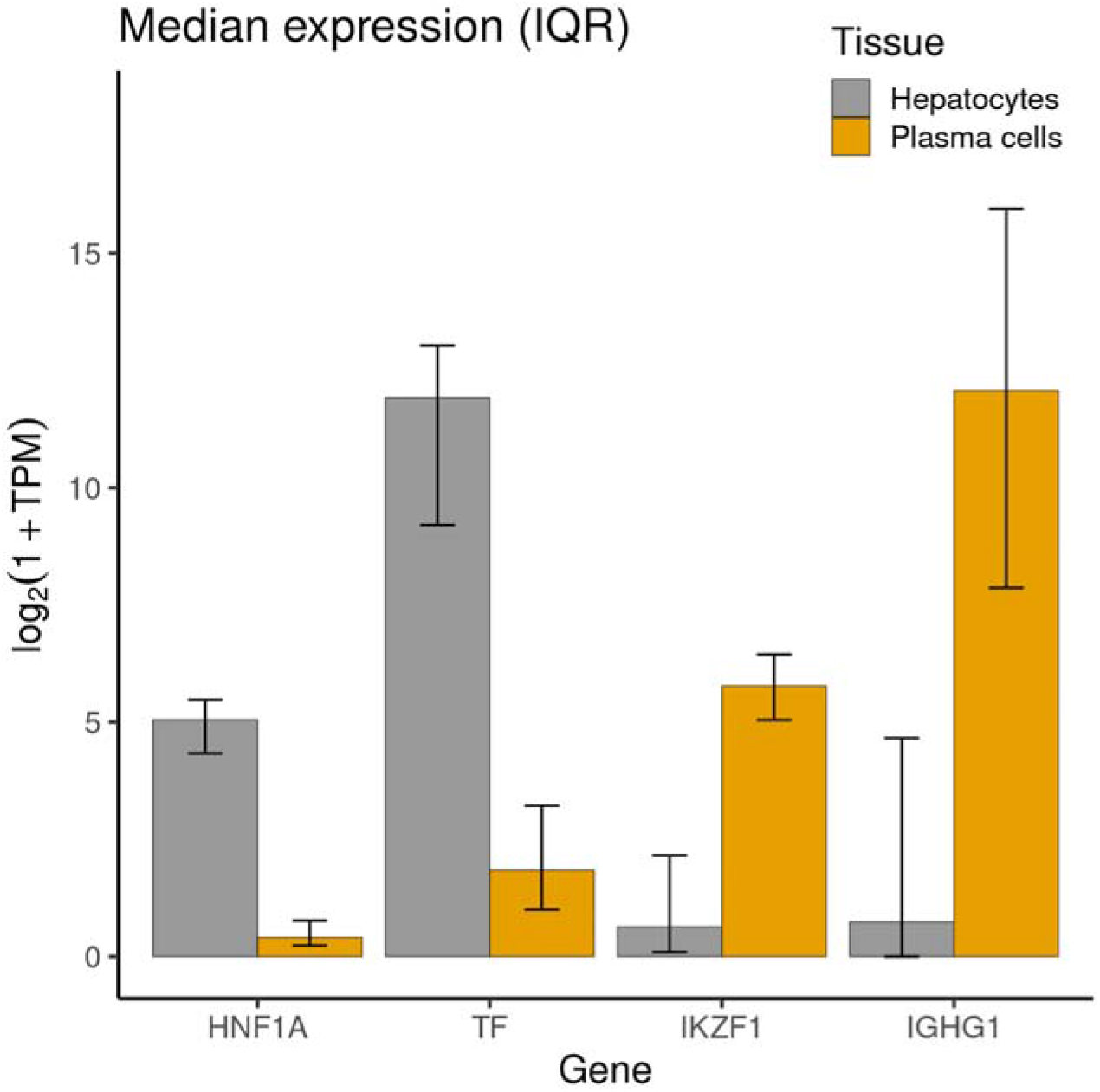
Expression of *HNF1A* and *IKZF1* in main tissues for transferrin and IgG proteins synthesis. Log_2_ of median transcripts per million (TPM) and interquartile ranges are reported for *TF* (encoding transferrin protein), *HNF1A, IGHG1* (encoding the constant region of immunoglobulin heavy chains) and *IKZF1* genes in plasma cells and hepatocytes. Gene expression data was obtained from the ARCHS4 portal^50^.

## Discussion

The post-translational modifications (PTMs) are essential mechanisms that dynamically regulate a large portion of cellular events by altering the structure and properties of proteins^1^. Similarly to other PTMs, genetic regulation of protein N-glycosylation has not been extensively investigated. Here, we performed genome-wide association meta-analysis of glycosylation of two proteins - transferrin and IgG - and compared how glycosylation of the two different proteins is genetically regulated. In the first GWAS of the transferrin N-glycome, (N=1890), we identified 10 significantly associated loci, three of which (near *TF, FOXI1* and *MSR1*) were never previously associated with the glycome of any protein. The other seven have been previously associated with glycosylation of total plasma proteins and/or IgG (Supplementary Table 2). The total plasma glycome quantifies the glycome of all proteins in plasma, but without information on which glycan was bound to which protein. Given that IgG and transferrin are among the most abundant plasma glycoproteins^12^, an overlap in genetic control of transferrin and IgG N-glycomes with that of total plasma proteins is to be expected. Sharapov et al.^28^ previously indicated that some of the genomic loci associated with the plasma glycome overlap with loci associated with IgG N-glycosylation. The present work suggests that the *MGAT5, ST3GAL4*, and *B3GAT1* loci, that were also observed in the total plasma protein GWAS, might be capturing a signal within plasma protein glycosylation that comes mainly from transferrin N-glycosylation.

We then compared the genetic architecture underlying glycosylation of transferrin and IgG proteins. Using the GWAS from this study we showed that there are both protein-specific and shared genetic loci. Looking specifically at glycosyltransferase enzymes, the main “drivers” of this post-translational modification, that catalyse the transfer of saccharide moieties from a donor to an acceptor molecule, *MGAT5, ST3GAL4*, and *B3GAT1* were only associated with transferrin while *ST6GAL1* and *MGAT3* were only associated with glycosylation of IgG. On the other hand, two fucosyltransferase genes, *FUT8* and *FUT6*, were associated with both proteins. Even though the genes encoding these enzymes were associated with glycosylation of both proteins, using Approximate Bayes Factor colocalisation analysis, we showed that associations with transferrin and IgG N-glycosylation at these genomic regions is driven by independent underlying causal variants, where one variant regulates fucosylation of transferrin and the other of IgG. Our results suggest that while the same fucosyltransferase enzymes are involved in N-glycosylation of both transferrin and IgG proteins, the process is independently regulated by protein-specific causal variants.

There are at least two mechanisms that could explain how different variants in an enzyme-coding gene could have distinct effects on two different substrates. If the two variants were in the coding region of the gene and affected the amino-acid sequence of the enzyme, they could affect the enzyme’s specificity for binding each protein. However, none of the sentinel variants in the *FUT8* and *FUT6* loci were in strong linkage disequilibrium (LD) with coding variants from the enzymes’ active sites, suggesting that this is likely not the mechanism of regulation of fucosylation of the two proteins. In addition, overall, SNPs associated with transferrin glycosylation predominantly mapped to regulatory rather than coding regions of the genome (Supplementary Table 5).

The other hypothesis is that these two variants affect the expression of enzymes in different tissues. In common with all other antibodies, most of IgG found in blood plasma is produced by bone marrow plasma cells, the fully differentiated form of B-cells^49^. The transferrin found in blood plasma is mostly produced by liver hepatocytes^48^. In addition, the glycomes of the two proteins were also associated with different transcription factor genes, namely, variants in *IKZF1* region were associated with IgG glycosylation, and variants in *HNF1A* region with transferrin glycosylation. IKZF1, a transcription factor predominantly expressed in immune cells and tissues, has been functionally validated as a regulator of IgG core fucosylation: IKZF1 binds to regulatory regions of *FUT8* and, in turn, knockdown of *IKZF1* results in increased expression of *FUT8* and increased core fucosylation of IgG^30^. On the other hand, we showed that transferrin glycosylation-associated variants in the *FUT8* region might affect the binding of HNF1A, a transcription factor predominantly expressed in the liver. *HNF1A* has already been shown to regulate the expression of *FUT8* and *FUT6* and affects fucosylation of total plasma proteins^27^. Overall, we hypothesise that the two different causal variants affect the binding of different transcription factors in different tissues and therefore regulate the glycosylation of the two plasma proteins in a tissue-specific manner.

In addition to HNF1A, *FUT8* glycosylation-associated variants might also be affecting the binding of the FOXI1 transcription factor. However, unlike HNF1A, possible involvement of FOXI1 in the regulation of the transferrin fucosylation is to date unknown and would require functional validation. We also found that HNF1A binding could also be affected by associated variants in the *TF* and *ST3GAL4* genes. While these relationships were hitherto undocumented and need further supporting evidence, they may suggest that HNF1A might regulate multiple genes associated with transferrin N-glycosylation.

Finally, our findings are not only relevant for unravelling the genetic mechanisms behind N-glycosylation PTM but also contribute to understanding changes in N-glycan patterns involved in disease. The most strongly N-glycosylation-associated variant for the *TF* gene, rs6785596, was suggested by McClain et al.^36^ to regulate *TF* expression in adipose tissue (also evident in GTEx v7) and consequently modulating insulin sensitivity. Excessive body iron stores represent a risk factor for decreased insulin sensitivity and diabetes^51^. McClain et al.^36^ argue that genetic downregulation of *TF* expression in adipocytes has functional consequences for these cells′ iron homeostasis and is sufficient to cause insulin resistance in humans and in a cell culture model. However, this SNP has so far not been associated with diabetes or diabetes-related traits, suggesting that this relationship needs to be explored further. Moreoever, while *TF* variant rs6785596 is not associated with transferrin protein levels (pQTL), we can consider it as an example of a “cis-glyQTL”: a genomic locus that explains variation in glycosylation levels and is local to the gene encoding the protein being glycosylated. Similar was observed for IgG glycosylation, where associated variants were mapping to the *IGH* locus^32^, a genetic region encoding the heavy chain of immunoglobulin G. In addition, glycosylation SNPs in *NXPE1/NXPE4* locus were pleiotropic with ulcerative colitis, a disease with abberant glycosylation patterns.

In conclusion, by performing the first GWAS of the plasma transferrin N-glycome and comparing it with that of the IgG N-glycome, we were for the first time able to describe similarities and differences in the genetic regulation of post-translational modification of two different proteins. When focusing on glycosyltransferases, main enzymes of this PTM, we showed that there are both associations specific to each protein, but also those that are shared in glycosylation of the two proteins. For the shared associations, we showed that fucosylation of transferrin and IgG are regulated by independent, protein-specific variants in the *FUT8* and *FUT6* genes. In the *FUT8* region these variants are likely to regulate fucosylation of transferrin and IgG in a tissue-specific manner, acting through tissue-specific transcription factors. Additional studies, with larger sample sizes and focusing on other non-IgG proteins, will be necessary to further unravel the genetic architecture of the N-glycosylation and other PTMs and to understand their relationship with human diseases and complex traits.

## Materials and Methods

### Population cohorts

The CROATIA-Korcula isolated population cohort includes samples of blood DNA, plasma and serum, anthropometric and physical measurements, information related to general health, medical history, lifestyle, and diet for ~3000 residents of the Croatian island of Korčula^52^. Written informed consent was given and the study was approved by the Ethics Committee of the Medical School, University of Split (approval id: 2181-198-03-04/10-11-0008). The Viking Health Study - Shetland (VIKING) is a family-based, cross-sectional study that seeks to identify genetic factors influencing cardiovascular and other disease risk in the population isolate of the Shetland Isles in northern Scotland^53^. Genetic diversity in this population is decreased compared to mainland Scotland, consistent with the high levels of endogamy. 2105 participants were recruited between 2013 and 2015, most having at least three grandparents from Shetland. Fasting blood samples were collected and many health-related phenotypes and environmental exposures were measured in each individual. All participants gave written informed consent and the study was approved by the South East Scotland Research Ethics Committee, NHS Lothian (reference: 12/SS/0151). Details of cohort-specific demographics, genotyping, quality control, and imputation performed before GWAS can be found in Supplementary Table 11.

### Phenotypic data

#### Transferrin and IgG N-glycome quantification

Transferrin and IgG N-glycome quantification for CROATIA-Korcula and VIKING samples was performed at Genos Glycobiology Laboratory. Isolation of the protein of interest and N-glycan quantification is described in more detail in Supplementary Materials and Methods for transferrin and by Trbojević-Akmačić et al.^54^ for IgG. Briefly, proteins were first isolated from blood plasma (IgG depleted blood plasma in the case of transferrin) using affinity chromatography binding respectively to anti-transferrin antibodies plates for transferrin and protein G plates for IgG. The proteins isolation step was followed by enzymatic release and labelling of N-glycans with 2-AB (2-aminobenzamide) fluorescent dye. N-glycans were then separated and quantified by hydrophilic interaction ultra-high-performance liquid chromatography (HILIC-UHPLC). As a result, transferrin and IgG samples were separated into 35 (transferrin: TfGP1-TfGP35) and 24 (IgG: GP1-GP24) chromatographic peaks. It is worth noting that there is no correspondence structure-wise between transferrin TfGP and IgG GP traits labelled with the same number.

#### Normalisation and batch correction of glycan traits

Prior to genetic analysis, raw N-glycan UHPLC data was normalised and batch corrected to reduce the experimental variation in measurements. Total area normalisation was performed by dividing the area of each chromatographic peak (35 for transferrin, 24 for IgG) by the total area of the corresponding chromatogram. Due to the multiplicative nature of measurement error and right-skewness of glycan data, normalised glycan measurements were log10-transformed. Batch correction was then performed using the empirical Bayes approach implemented in the “ComBat” function of the “sva” R package^55^, modelling the technical source of variation (96-well plate number) as batch covariate. Batch corrected measurements where then exponentiated back to the original scale.

### Genome-wide association analysis

Genome-wide association analyses (GWAS) were performed in the two cohorts of European descent, CROATIA-Korcula and VIKING. Associations with 35 transferrin N-glycan traits were performed in 948 samples from CROATIA-Korcula and 959 samples from VIKING. Associations with 24 IgG N-glycan traits were performed in 951 samples from CROATIA-Korcula and 1086 samples from VIKING. The sample size of the same cohort differs between transferrin and IgG due to the different number of samples successfully measured for each protein. Prior to GWAS, each glycan trait was rank transformed to normal distribution using the “rntransform” function from the “GenABEL” R package^56^ and then adjusted for age and sex, as fixed effects, and relatedness (estimated as the kinship matrix calculated from genotyped data) as random effect in a linear mixed model, calculated using the “polygenic” function from the “GenABEL” R package^56^. Residuals of covariate and relatedness correction were tested for association with HRC (Haplotype Reference Consortium) imputed SNP dosages using the RegScan v. 0.5 software^57^, applying an additive genetic model of association.

### Meta-analysis

#### Meta-analysis

Prior to meta-analysis the following quality control was performed on cohort-level GWAS summary statistics. We removed all SNPs with a difference in allele frequency between the two cohorts higher than +/- 0.3, as well as variants showing a minor allele count (MAC) lower or equal to 6. Cohort-level GWAS were then meta-analysed (N=1890 for transferrin and N=2020 for IgG N-glycans, for ~12 million SNPs) using METAL software^58^, applying the fixed effect inverse-variance method, followed by genomic control correction. Mean genomic control inflation factor (λ_GC_) was 0.997 (range 0.982-1.011) for transferrin N-glycans and 0.995 (range 0.981-1.008) for IgG N-glycans meta-analysis, showing that the confounding effects of family structure were correctly accounted for.

#### Multiple test correction

The standard genome-wide significance threshold was Bonferroni corrected for the number of N-glycan traits analysed: variants were considered statistically significant if their p-value was lower than 5×10^-8^/35 = 1.43×10^-9^ for transferrin and 5×10^-8^/24= 2.08×10^-9^ for IgG N-glycan traits.

#### Locus definition

We used a positional approach to define genomic regions significantly associated with transferrin N-glycan traits, following the procedure adopted by Sharapov et al.^28^ For each glycan trait, we grouped all genetic variants located within a 500 kb window (+/- 250 kb) from the sentinel SNP in the same locus. To obtain a unique list of loci that are independent of the specific glycan trait, we then merged this list of sentinel SNP-glycan trait pairs for all 35 glycan traits and applied a similar procedure - all SNP-glycan trait pairs within a 1000 kb window (+/-500 kb from sentinel SNP) were grouped in the same locus, resulting in a unique list of sentinel SNP-top glycan trait pairs, summarising the genomic regions most strongly associated with N-glycans across all traits. For all sentinel SNP-top glycan trait pairs, regional association plots were created with LocusZoom^59^ and visually checked - in case of overlapping patterns of association, only the sentinel SNP-top glycan trait pair showing the lowest p-value was selected as a locus representative.

### Transferrin N-glycan traits post-meta-analysis follow-up

The meta-analysis follow-up analyses were performed only for the transferrin N-glycans meta-analysis, since genetic regulation of IgG N-glycosylation has already been explored in a larger, IgG-specific study^30^ and is beyond the scope of the present work.

#### Conditional analysis and phenotypic variance explained

To capture the overall contribution to phenotypic variation at each genomic region and identify secondary association signals at a locus, we performed approximate conditional analysis using the GCTA-COJO^39^ stepwise model selection, “cojo-slct”, with the transferrin N-glycan metaanalysis summary statistics and genotypes of 10,000 unrelated individuals of white British ancestry from UK Biobank^60^ as independent LD reference panel. Collinearity was restricted to 0.9 and the p-value threshold was set to 1.43×10^-9^. Reported joint p-values were then adjusted by the genomic control method^61^. The list of samples for the independent LD reference panel was created with R 3.6.0, while the panel itself was generated using Plink 2.0^62^. After samples extraction from the UK Biobank full dataset, SNP deduplication was performed both by position (removing all SNPs not carrying a unique position on the chromosome) and marker name (--rm-dup exclude-all function). The proportion of variance in phenotype (*Y*) explained by sentinel SNPs at each transferrin N-glycans associated locus was calculated with the following formula

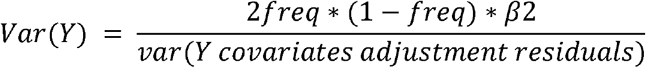

#### Gene prioritisation

For all genome-wide significant loci we suggested plausible candidate genes combining different evidence, namely evaluating biological role in the context of protein N-glycosylation of genes nearest to sentinel variants (positional mapping), assessing pleiotropy of sentinel variants with gene expression (expression quantitative trait loci, eQTL) or investigating associated variant’s predicted effects on the protein sequence or on putative transcription factor binding sites (TFBSs). Positional gene mapping was performed using FUMA v1.3.5e SNP2GENE function^63^. Genes having a clear biological link to protein N-glycosylation (e.g. genes coding for enzymes involved in biochemical pathway of protein glycosylation) and genes previously associated with IgG and/or total blood plasma proteins N-glycome were given a priority. The overlap of independent significant SNPs identified by COJO with eQTL was investigated using PhenoScanner v1.1 database^45^, taking into account significant genetic association (p-value < 5×10 ^–8^) at the same or strongly (LD r^2^ > 0.8) linked SNPs in populations of European ancestry. The Ensembl Variant Effect Predictor (VEP v 97) tool^42^ was used to determine putative functional effect and impact on a transcript or protein of independent significant SNPs and their strongly (LD r^2^ > 0.8) linked SNPs in populations of European ancestry. Among genes prioritised so far, two were transcription factors (i.e. *HNF1A* and *FOXI1*), while the remaining were non transcription factor protein-coding genes (i.e. *MGAT5, TF, MSR1, NXPE1/NXPE4, ST3GAL4, B3GAT1, FUT8* and *FUT6*). Using the Regulatory sequence analysis tools (RSAT) program *malπx-scan^44^*, we applied a pattern-matching procedure to search for sequences recognized as binding sites for *HNF1A* and *FOXI1* transcription factors in associated regions of the other 8 prioritised genes. Position-specific scoring matrices (PSSMs), representing the frequency of each nucleotide at each position of the transcription factor motif, were downloaded for *HNF1A* and *FOXI1* from the JASPAR^64^ database. For each of the 8 genomic regions explored for possible transcription factor binding sites, we included the most strongly associated SNP and a 60 bp surrounding sequence (30 bp either side of the sentinel SNP). The significance threshold was set to the p-value ≤ 0.003, Bonferroni corrected for 16 tests performed (8 putative transcription factor binding sites tested for 2 transcription factors).

#### Overlap and colocalization analysis with gene expression levels and complex traits

The PhenoScanner v1.1 database^45^ was used to investigate the overlap of significant transferrin glycosylation SNPs with gene expression levels and complex human traits. As previously described, we considered traits with genome-wide significant association (p-value < 5×10^−8^) at the same or strongly (LD r^2^ > 0.8) linked SNPs in populations of European ancestry. We then used Summary data-based Mendelian Randomization (SMR) analysis followed by the Heterogeneity in Dependent Instruments (HEIDI) test^41^ to assess whether overlapping expression and complex traits, identified by PhenoScanner, were also colocalising with transferrin glycosylation (TfGP) traits. The SMR test indicates whether two traits are associated with the same locus, and HEIDI test specifies whether both traits are affected by the same underlying functional SNP. Each of 10 sentinel SNPs – TfGP pair (Table 1) was used for SMR/HEIDI analysis with gene expression levels and several complex traits. Summary statistics for gene expression levels in tissues/cell types were obtained from the Blood eQTL study^65^ (http://cnsgenomics.com/software/smr/#eQTLsummarydata), the CEDAR project^66^ (http://cedar-web.giga.ulg.ac.be/), and the GTEx project version 7^67^ (https://gtexportal.org). Summary statistics for complex traits were obtained from various resources. In total, we used data for 3 tissues/cell types: CD19+ B lymphocytes (CEDAR), GTEx liver (GTEx) and peripheral blood (the Blood eQTL study) and 8 complex traits. Full list of GWAS collections, tissues and complex traits see in Supplementary Table 12. SMR/HEIDI analysis was performed according to the protocol described by Zhu et al.^41^ We used sets of SNPs having the following properties: (1) being located within ± 250 kb from the sentinel SNPs identified in the present study; (2) being present in both the primary GWAS and eQTL data/GWAS for the complex trait; (3) having MAF ≥ 0.03 in both datasets; (4) having squared Z-test value ≥ 10 in the primary GWAS. Those SNPs that met criteria (1), (2), (3), (4), had the lowest P-value in the primary GWAS and were in high LD (r^2^□>□0.8) with the sentinel SNPs were used as instrumental variables to elucidate the relationship between gene expression/disease and TfGP (we define them as “top SNPs”). It should be noted that SMR/HEIDI analysis does not identify a causative SNP affecting both traits. It can be either the top SNP or any other SNP in strong LD. After defining the set of eligible SNPs for each locus, we made the “target” and “rejected” SNP sets and added the top SNP to the “target” set. Then we performed the following iterative procedure of SNP filtration: if the SNP from the eligible SNP set with the lowest PSMR had r^2^ > 0.9 with any SNP in the “target” SNP set, it was added to the “rejected” set; otherwise, it was added to the “target” set. The procedure was repeated until eligible SNP set was exhausted, or the “target” set had 20 SNPs. If we were unable to select three or more SNPs, the HEIDI test was not conducted. HEIDI statistics was calculated as 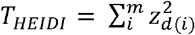, where m is the number of SNPs selected for analysis,

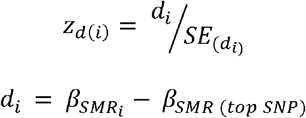

The results of the SMR test were considered statistically significant if PSMR < 1.7 × 10^-4^ (0.05/302, where 302 is a total number of tests corresponding to analyzed loci and gene expression/disease traits). Inference of whether a functional variant may be shared between the TfGP and gene expression/disease were made based on the HEIDI test: P_HEIDI_ ≥ 0.001 (possibly shared), and P_HEIDI_ < 0.001 (sharing is unlikely).

### Colocalisation analysis for transferrin and IgG N-glycan traits

The *FUT8* and *FUT6* genomic regions were significantly associated with both transferrin and IgG N-glycans. To investigate a possible overlap in genetic control of glycosylation between the two proteins, we used the approximate Bayes factor colocalisation analysis, developed by Giambartolomei et al.^68^ and implemented in “coloc” R package, followed by pairwise conditional and colocalization analysis (PwCoCo)^47^ in case of multiple independent variants contributing to the trait variation. A posterior probability (PP) > 80% was considered as robust evidence supporting the tested hypothesis.

Overview of the overall procedure can be seen in Supplementary Figure 1. First, we assessed whether for one protein all glycans that are associated with the same genomic region (p-value ≤ 5×10^-8^) are regulated by the same underlying variants. For each protein (i.e. transferrin and IgG) and each genomic region (i.e. *FUT8* and *FUT6*), we tested separately the group of glycans carrying only one independent association signal at locus and the group of glycan traits showing multiple independent signals of association (Supplementary figure 1). Pairs of glycan traits obtaining a PP.H4 > 80% (suggestive of colocalisation) were pooled in the same colocalisation group, following the principle that if trait A colocalises with trait B and trait B colocalises with trait C, thus also trait A and trait C colocalise. For each within-protein colocalisation group identified, the glycan trait with the lowest p-value was selected as group representative and carried on to the next step, where traits with single and multiple independent associations for each protein were tested for colocalisation. Similar to previous steps, glycan traits were grouped together on the basis of their colocalisation analysis results and the lowest p-value representative was chosen for the next step, where finally representative transferrin and IgG glycans were tested for between-protein colocalisation.

For glycan traits with multiple independent association signals and lacking strong evidence for colocalisation, we applied PwCoCo^47^ approach. Briefly, the PwCoCo approach tests not only the traits’ full, complete GWAS association statistics for colocalisation, but also summary statistics conditioned for the top primary association, testing whether any of the underlying causal variants between traits colocalise. For example, assuming that each trait is carrying two conditionally independent association signals in the tested region, colocalisation analysis will be conducted between both full and conditioned association statistics (conditioned for each independent variable), for a total of nine pairwise combinations. Secondary association signals at *FUT8* and *FUT6* loci for both transferrin and IgG N-glycans were assessed using GCTA-COJO approximate conditional analysis stepwise model selection^39^ and an LD reference panel of 10,000 unrelated, white British ancestry individuals from UK Biobank^60^. We then performed the association analysis conditional on identified secondary association signals at *FUT8* and *FUT6* loci using GCTA-COJO^39^ “cojo-cond” and the same 10,000 UK Biobank samples LD reference panel, with 5×10^-8^ p-value threshold and used those for pairwise colocalisation analyses.

### Expression of N-glycome associated genes in transferrin and IgG relevant tissues

Gene expression data for *TF, IGHG1, HNF1A* and *IKZF1*, expressed in gene counts, for hepatocytes (529 samples) and plasma cells (648 samples) was obtained from ARCHS4 portal^50^. Samples with total number of gene counts less than 5,000,000 were filtered out. 513 hepatocyte and 53 plasma cell samples were undergone further analysis. Gene counts were scaled to transcripts per million (TPM) and log2(1+TPM) transformed.

## Supporting information

Supplementary Methods, Reults and Figures

## Acknowledgements

The CROATIA_Korcula study was funded by grants from the MRC (United Kingdom), European Commission Framework 6 project EUROSPAN (contract number LSHG-CT-2006-018947), Croatian Science Foundation (grant 8875), and the Republic of Croatia Ministry of Science, Education and Sports (216-1080315-0302). Genotyping was performed in the Genetics Core of the Clinical Research Facility, University of Edinburgh.We would like to acknowledge all the staff of several institutions in Croatia that supported the CROATIA_ Korcula fieldwork, including, but not limited to, the University of Split and Zagreb Medical Schools, Institute for Anthropological Research in Zagreb, and the Croatian Institute for Public Health in Split. The Viking Health Study – Shetland (VIKING) was supported by the MRC Human Genetics Unit quinquennial programme grant “QTL in Health and Disease”. DNA extractions and genotyping were performed at the Edinburgh Clinical Research Facility, University of Edinburgh. We would like to acknowledge the invaluable contributions of the research nurses in Shetland, the administrative team in Edinburgh and the people of Shetland. We acknowledge support from the MRC Human Genetics Unit programme grant, “Quantitative traits in health and disease” (U. MC_UU_00007/10).

A.L. was funded by the European Union’s Horizon 2020 research and innovation program IMforFUTURE, under H2020-MSCA-ITN grant agreement number 721815. The work of L.K. was supported by an RCUK Innovation Fellowship from the National Productivity Investment Fund (MR/R026408/1). The work of Y.S.A. was supported by a grant from the Russian Science Foundation (RSF) No. 19-15-00115. The work of J.F.W. and C.H. was supported by an MRC University Unit Programme Grant MC_UU_00007/10 “QTL in Health and Disease”.

## Author contributions

A.L.: Data analysis and interpretation, Visualization, Writing—Original draft preparation, Writing—Review and editing. I.T.-A.: Quantification of transferrin and IgG N-glycans, Data interpretation, Writing—Original draft preparation, Writing—Review and editing. P.N.: Supervision, Data interpretation, Writing—Review and editing. Y.T.: Data analysis and interpretation, Writing—Original draft preparation. S.Z.S.: Visualization, Writing—Original draft preparation. F.V.: Glycan data quality control. O.P.: Genomic and demographic data provider for CROATIA-Korcula cohort. C.H.: Genomic and demographic data provider for CROATIA-Korcula cohort. T.P.: Quantification of transferrin and IgG N-glycans. M.V.: Quantification of transferrin and IgG N-glycans. Y.S.A.: Writing—Review and editing. G.L.: Glycan data provider for CROATIA-Korcula and VIKING cohorts, Writing—Review and editing. J.F.W.: Conceptualization, Genomic and demographic data provider for VIKING cohort, Supervision, Data interpretation, Writing—Original draft preparation, Writing— Review and editing. L.K.: Conceptualization, Supervision, Data interpretation, Writing— Original draft preparation, Writing—Review and editing.

## Competing interests

G.L. is the founder and owner of Genos Ltd, a private research organization that specializes in high-throughput glycomic analysis and has several patents in this field. I.T.-A., F.V., T.P., and M.V. are employees of Genos Ltd. Y.S.A. is a founder and a co-owner of PolyOmica and PolyKnomics, private organizations providing services, research and development in the field of computational and statistical genomics.

