## Supplementary Methods, Reults and Figures for "Same role but different actors: genetic regulation of post-translational modification of two distinct proteins"

**Supplementary Materials and Methods**

**Transferrin isolation**

Flowthrough during IgG purification was collected for immediate subsequent Tf isolation using previously developed preconditioned CIMac-@Tf 96-well monolithic plate^1^. Unbound proteins during Tf isolation were washed away with 1x PBS (0.25 mol L^-1^ NaCl), pH 7.4. Bound Tf was eluted with 0.7 mL of 0.1 mol L^-1^ formic acid pH 3.0 (pH adjusted with 25 % ammonia solution, Merck) and immediately neutralized with 1 mol L^-1^ ammonium hydrogencarbonate (Sigma-Aldrich) to pH 7.0. Monolithic plate was regenerated and stored at 4 °C until the next isolation. Each elution fraction (300 μL) was dried in a vacuum centrifuge (Thermo Scientific) and stored at -20 °C until subsequent N-glycan release.

**N-glycan release and fluorescent labeling**

Dried Tf eluates were denatured with 30 μl of 13.3 g L^-1^ sodium dodecyl sulfate (SDS, Invitrogen) and by incubation at 65 °C for 10 min. After cooling down to room temperature for 30 min, 10 μl of 4 % (v/v) Igepal CA-630 (Sigma-Aldrich) was added and the mixture was shaken for 15 min on a plate shaker. N-glycans were released after the addition of 10 μL of 5x PBS and 1.2 U of PNGase F (Promega) by incubation at 37 °C for 18 hours. Released N-glycans were labeled with 2-aminobenzamide (2-AB, Sigma-Aldrich). The labeling mixture was freshly prepared by dissolving 2-AB and 2-methylpyridine borane complex (2-PB, Sigma-Aldrich) (final concentrations of 19.2 mg mL^-1^ and 44.8 mg mL^-1^, respectively) in the mixture of dimethyl sulfoxide (Sigma-Aldrich) and glacial acetic acid (Merck) (7:3). Labeling mixture (25 μL) was added to each sample and the plate was sealed using an adhesive seal. After 10 minutes of shaking, samples were incubated for 2 hours at 65 °C. Excess of reagents from previous steps was removed from the samples using hydrophilic interaction liquid chromatography solid phase extraction (HILIC-SPE). After free N-glycan labeling samples were cooled down to room temperature for 30 min and 700 μL of acetonitrile (previously cooled down to 4 °C) was added to each sample. The cleanup procedure was performed on a hydrophilic 0.2 μm AcroPrep GHP filter plate (Pall) using a vacuum manifold (Pall) at around 25 mm Hg. All wells of a GHP filter plate were prewashed with 200 μL of 70 % (v/v) ethanol in water, 200 μL of ultrapure water, and 200 μL of 96 % (v/v) acetonitrile in water (previously cooled down to 4 °C). Diluted samples were loaded to the GHP filter plate wells, and after short incubation subsequently washed with 5x 200 μL of 96 % (v/v) acetonitrile in water. The last washing step was followed by centrifugation at 164 *g* for 5 minutes. Glycans were eluted from the plate with 2x 90 μL of ultrapure water after 15 min shaking at room temperature and centrifugation at 164 *g* for 5 minutes in each step. Combined eluates of 2-AB labeled Tf N-glycans were stored at -20 °C until ultra-high-performance liquid chromatography (UHPLC) analysis.

**Glycan analysis by ultra-high-performance liquid chromatography**

Fluorescently labeled and purified Tf N-glycans were analyzed by UHPLC based on hydrophilic interactions (HILIC-UHPLC) and detected using excitation and emission wavelengths of 250 and 428 nm, respectively. Acquity UHPLC instrument (Waters) was under the control of Empower 3 software, build 3471 (Waters). Mobile phases were 100 mmol L^-1^ ammonium formate, pH 4.4 (solvent A) and acetonitrile (solvent B) and samples were maintained at 10 °C before injection. Tf 2-AB labeled N-glycans prepared in 75 % acetonitrile were separated on a Waters BEH Glycan column, 150 × 2.1 mm i.d., 1.7 μm BEH particles at 25 °C in a linear gradient of 30-47 % solvent A at a flow rate of 0.56 mL min^-1^ during a 23 minute analytical run.

The HILIC-UHPLC system was calibrated using a dextran ladder (external standard of hydrolyzed and 2-AB labeled glucose oligomers) according to which the retention times for the individual chromatographic peaks (representing the 2-AB labeled glycan) were converted to glucose units (GU). Data processing was performed using an automatic processing method with a traditional integration algorithm. Each Tf N-glycans chromatogram integrated into 35 peaks was manually corrected to maintain the same intervals of integration for all the samples. The amount of glycans in each chromatographic peak was expressed as a percentage of the total integrated area (% Area).

**Replication of transferrin N-glycans loci**

To assess robustness of our findings we used the VIKING cohort as replication cohort and CROATIA-Korcula as discovery cohort. Each significant sentinel SNP-top glycan trait pair from the discovery cohort was tested for associations in the replication cohort, with replication significance threshold set to the p-value ≤ 0.00625 (0.05/8, number of discovery cohort genome-wide significant loci). Where the SNP of interest was not available in the replication cohort, a proxy SNP in high linkage disequilibrium (r2 ≥ 0.8) was used instead. In addition to statistical significance, we also assessed if the direction of estimated effect was concordant between discovery and replication study.

**Supplementary Results**

**Discovery and replication transferrin N-glycans GWAS**

We performed GWAS of 35 ultra-high-performance liquid chromatography (UHPLC) measured transferrin N-glycan traits and Haplotype Reference Consortium (HRC) r1.1 imputed genetic data in two cohorts of European descent, CROATIA-Korcula (N=938) and VIKING (N=952). Overall, we identified 8 loci genome-wide significantly associated (p-value ≤ 1.43×10^-9^) with transferrin N-glycans in the CROATIA-Korcula discovery cohort (Supplementary Figure 5, top panel and Supplementary Table 13), 6 of which replicated in the VIKING cohort (p-value ≤ 0.00625) (Supplementary Figure 6 and Supplementary Table 13). Replicated loci contained genes encoding glycosyltransferases, enzymes directly involved in the biochemical pathway of N-glycosylation (i.e. *MGAT5*, *ST3GAL4*, *B3GAT1*, *FUT8* and *FUT6*) and the transferrin (*TF*) gene.

**Transferrin N-glycans shared genetic associations with complex traits and diseases**

For the shared associations between transferrin glycosylation from the *ST3GAL4* locus and LDL, total cholesterol levels and platelet-related traits; *HNF1A* and coronary artery disease, levels of C-reactive protein and of gamma-glutamyl transferase; *FUT6* and age-related macular degeneration (Supplementary Table 7a); the SMR p-value was not significant (Supplementary Table 7b), so the inference on pleiotropy could not be performed.

**Colocalisation analysis of transferrin and IgG glycan traits with multiple independent association signals at genomic region**

To investigate whether the same variant within the *FUT8* and *FUT6* loci is regulating glycosylation of both proteins, and, at the same time, account for the presence of multiple conditionally distinct association signals within the same locus, we applied the PwCoCo pipeline^2^, integrating Approximate Bayes Factor (ABF) colocalisation^3^ and conditional analyses (for details see Supplementary Figure 1). Briefly, to address the problem of multiple associations within a locus, this approach tests for colocalisation using not only the trait’s unconditioned GWAS association statistics, but also their conditioned ones, assessing if any of the independent associations colocalise^2^. We first tested for evidence of multiple SNPs independently contributing to IgG glycan levels at the *FUT6* and *FUT8* loci. While no secondary associated variants were observed in the *FUT6* locus using GCTA-COJO stepwise analysis, two independent variants are likely to contribute to variation in transferrin and IgG glycan traits, namely transferrin TfGP32 and IgG GP20, in the *FUT8* locus (Supplementary Table 2). In this case, colocalisation analyses were conducted between full unconditioned association statistics, association statistics conditioned by one association signal (i.e. transferrin TfGP20 conditioned on rs72716459 and IgG GP7 conditioned on rs8022094) and those conditioned by the other association signal (i.e. transferrin TfGP20 conditioned on rs2411815 and IgG GP7 conditioned on rs8006608), for a total of nine pairwise combinations. We obtained robust evidence against the colocalisation hypothesis for all tested traits, except for transferrin TfGP20 conditioned on rs72716459 and IgG GP7 unconditioned association statistics. In this case in fact it was not possible to strongly support either the hypothesis of different causal variants at the locus or trait colocalisation (PP.H3 = 46.82%, PP.H4 = 53.18%) and therefore whether this glycans pair share genetic architecture at this locus remains unclear (Supplementary Table 10, Supplementary Figure 2 and Supplementary Figure 3).

It is important to note that another transferrin glycosylation associated genetic region, harbouring *NXEP1* and *NXEP4* genes, was also associated with IgG glycosylation in Klaric et al^4^. Since the role of these genes is unknown and therefore interpretation of their role in glycosylation is not straightforward, we did not proceed with colocalisation analysis for this region.

**Supplementary Figures**

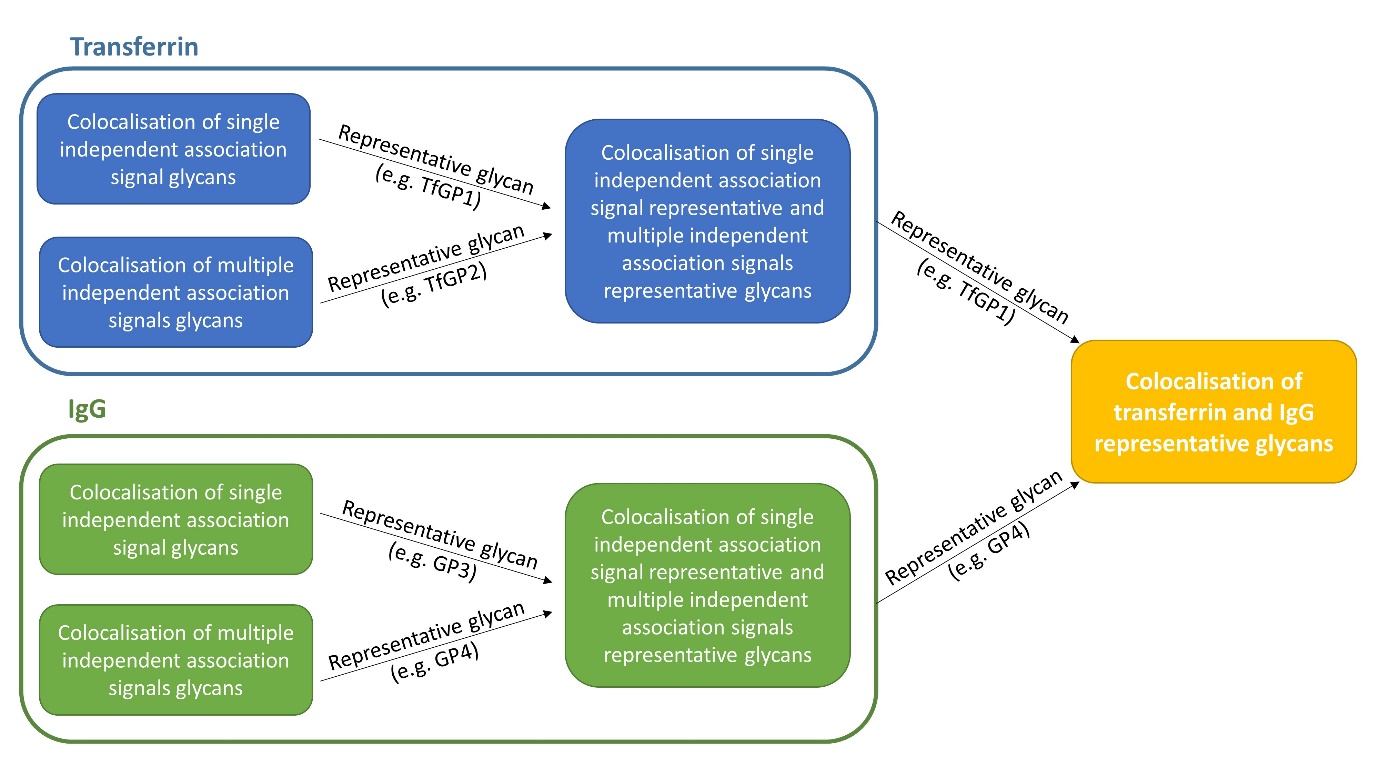

**Supplementary Figure 1. Workflow applied for transferrin and IgG glycan traits colocalisation analysis.** For each protein (i.e. transferrin and IgG) and each genomic region (i.e. *FUT8* and *FUT6*), the group of glycan traits showing multiple independent signals of association and, separately, the group of glycans carrying only one independent association signal at locus were pair-tested for traits colocalisation. Pairs of glycan traits obtaining an ABF posterior probability for hypothesis 4 (suggestive of colocalisation) > 80% were pooled in the same colocalisation group. For each colocalisation group identified, the glycan trait reporting the lowest p-value was selected as group representative and carried on to the next step, where traits with single and multiple independent associations for each protein were tested for colocalisation. Similar to previous steps, glycan traits were grouped together on the basis of their colocalisation analysis results and the lowest p-value representative was chosen for the next step, where finally representative transferrin and IgG glycans at each locus were tested for colocalisation.

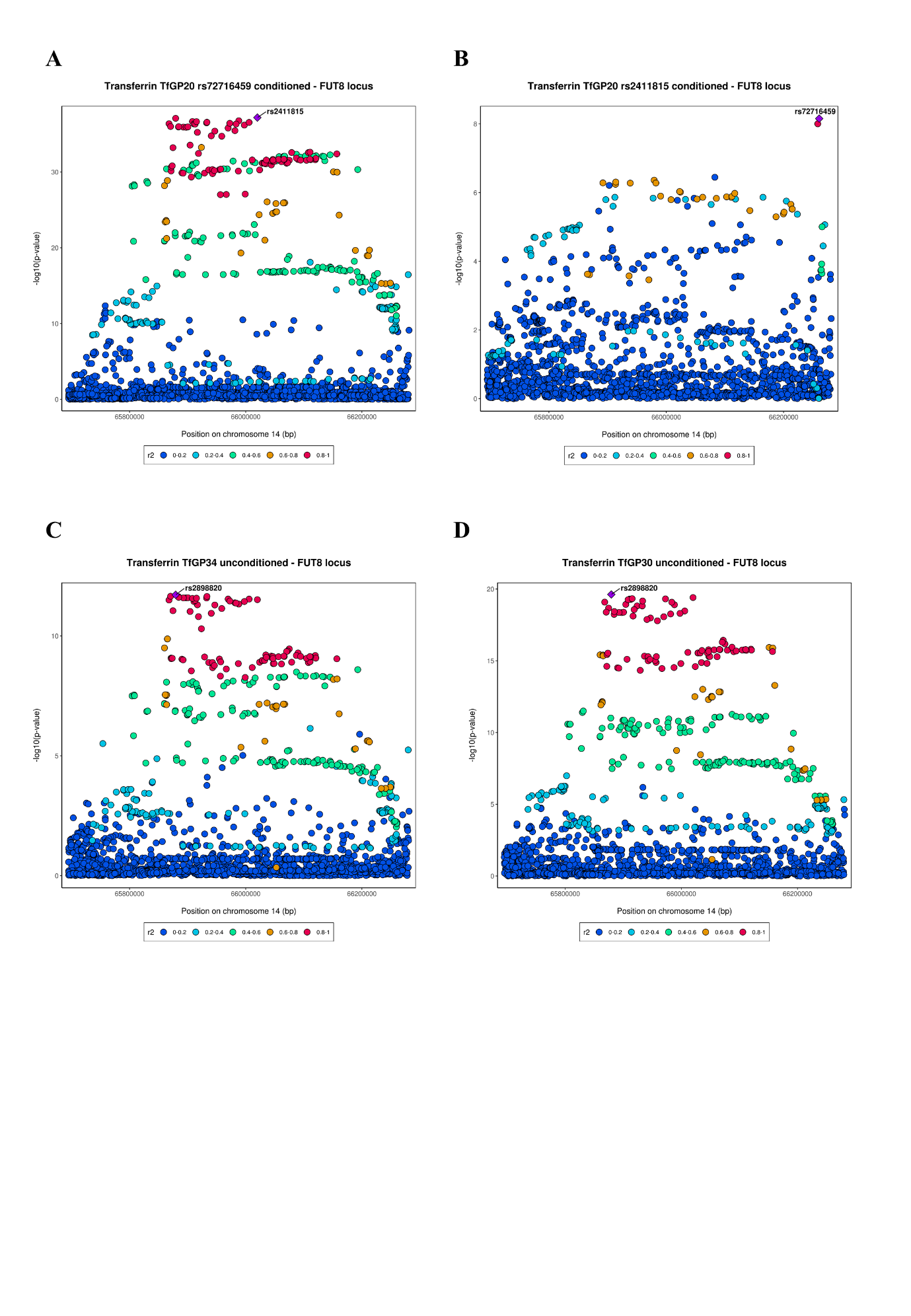

**Supplementary Figure 2. Local association patterns of transferrin N-glycans tested for trait colocalisation at *FUT8* locus.** Colocalisation analysis results for these glycan traits can be found at Supplementary Tables 9 (C and D) and S10 (A and B). Since TfGP20 also has an independent secondary association signal (see Supplementary Table 2), local association patterns are reported also for conditioned summary statistics.

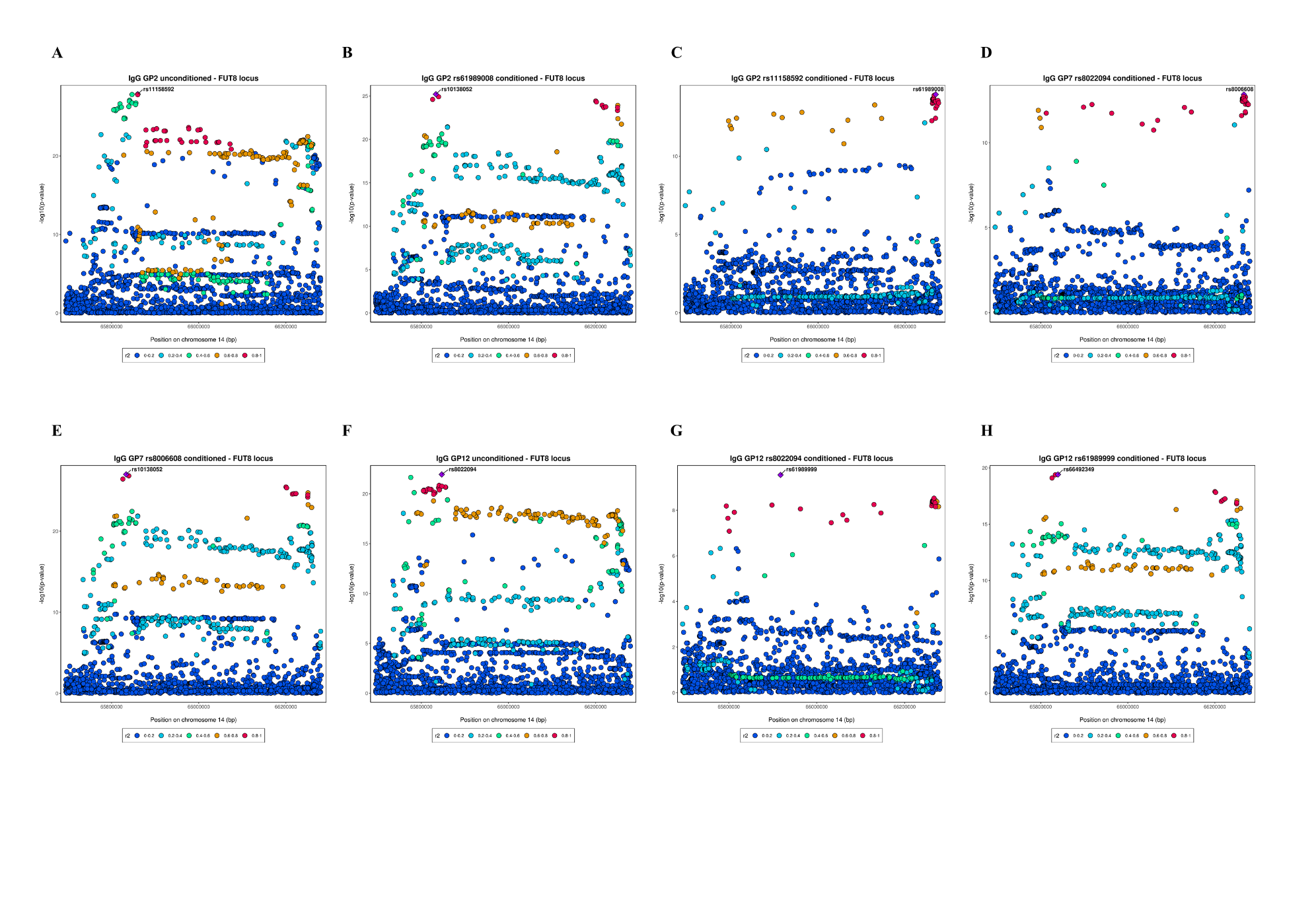

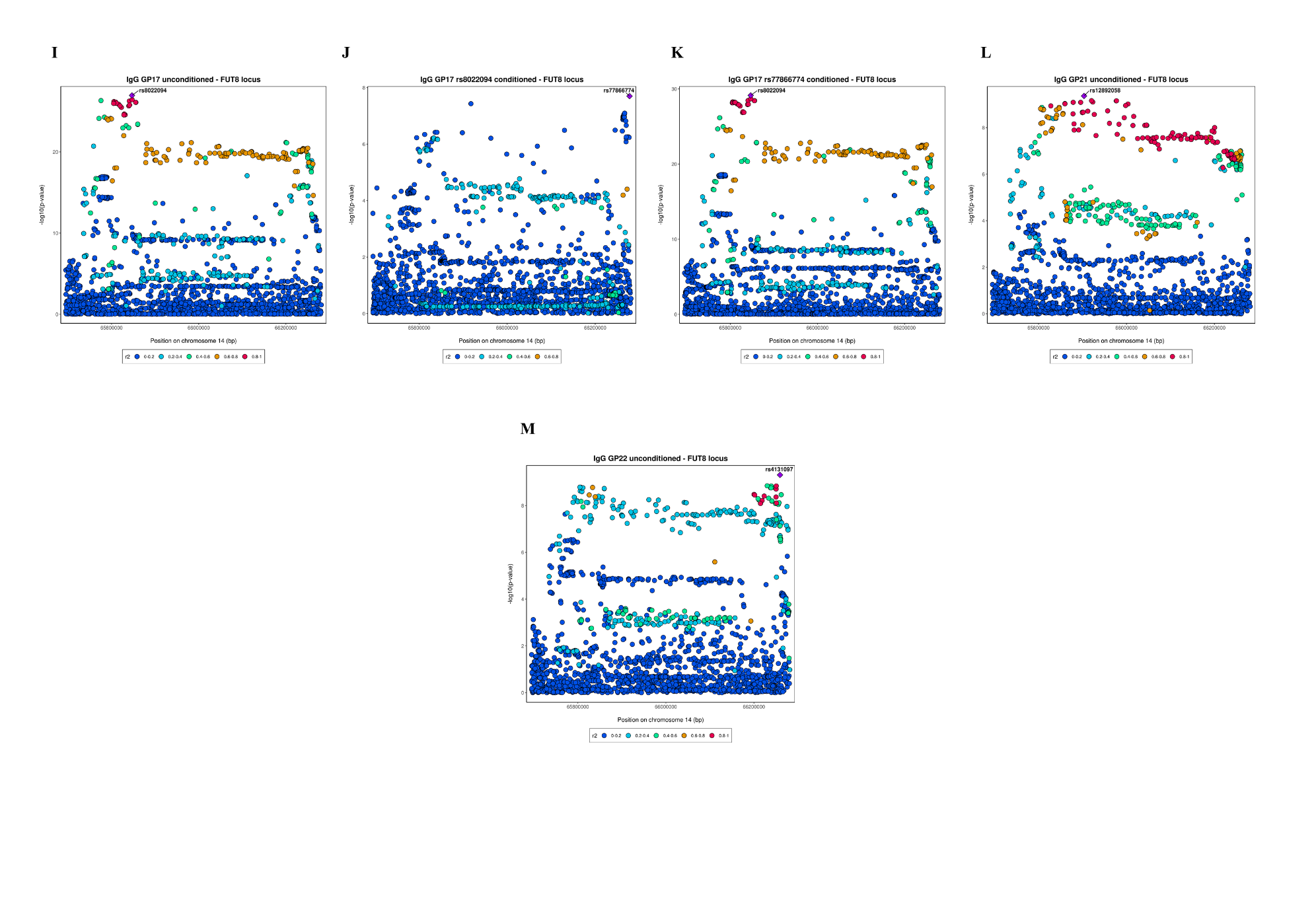
**Supplementary Figure 3. Local association patterns of IgG N-glycans tested for trait colocalisation at *FUT8* locus.** Colocalisation analysis results for these glycan traits can be found at Supplementary Table 9 (A, B, C, F, G, H, I, J, K, L, M and N) and S10 (D and E). Since GP7 also has an independent secondary association signal (see Supplementary Table 2), local association patterns are reported also for conditioned summary statistics.

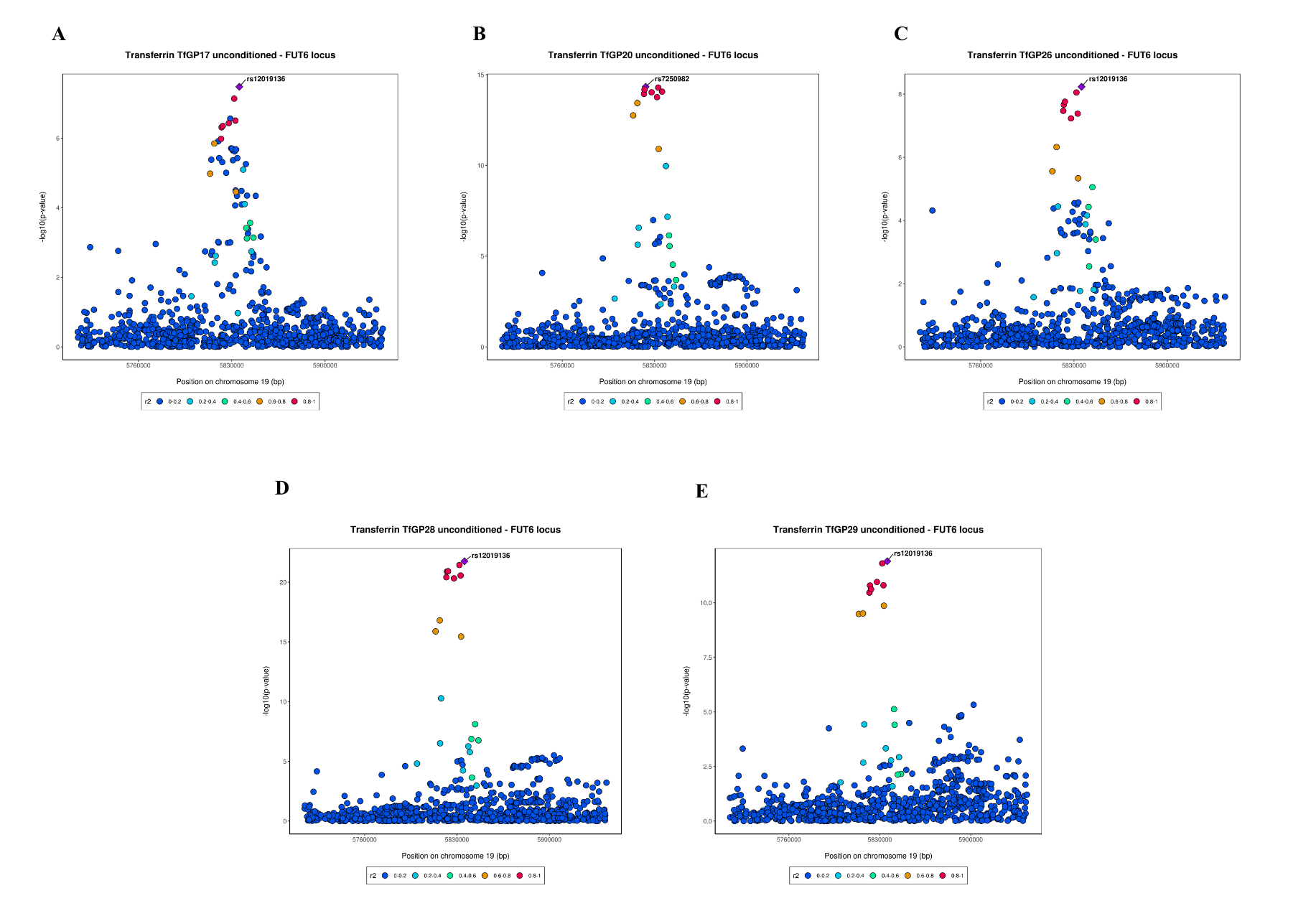
**Supplementary Figure 4. Local association patterns of transferrin N-glycans tested for trait colocalisation at *FUT6* locus.** Colocalisation analysis results for these glycan traits can be found at Supplementary Tables 9.

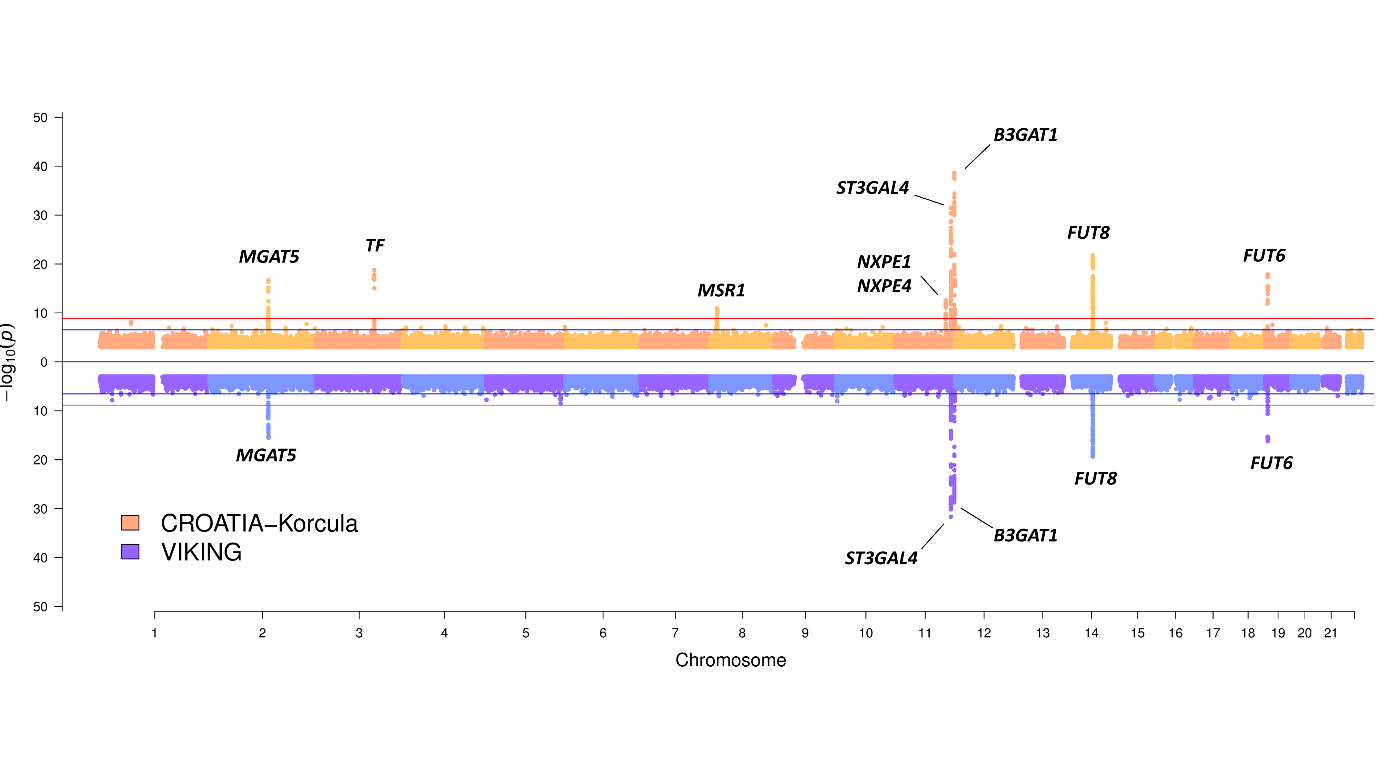
**Supplementary Figure 5. Transferrin N-glycome CROATIA-Korcula and VIKING cohorts GWAS summary Miami plot.** Miami plot pooling together individual cohort GWAS results obtained across all 35 transferrin glycan traits, at the top in orange for CROATIA-Korcula cohort and at the bottom in blue for VIKING cohort. For each SNP, the lowest p-value scored across the 35 traits is reported. The horizontal red line corresponds to the multiple testing corrected genome-wide significance threshold of 1.43×10^-9^. The horizontal blue line corresponds to the multiple tests corrected genome-wide suggestive threshold of 2.86×10^-7^. For simplicity, SNPs with p-value > 1×10^-3^ are not reported.

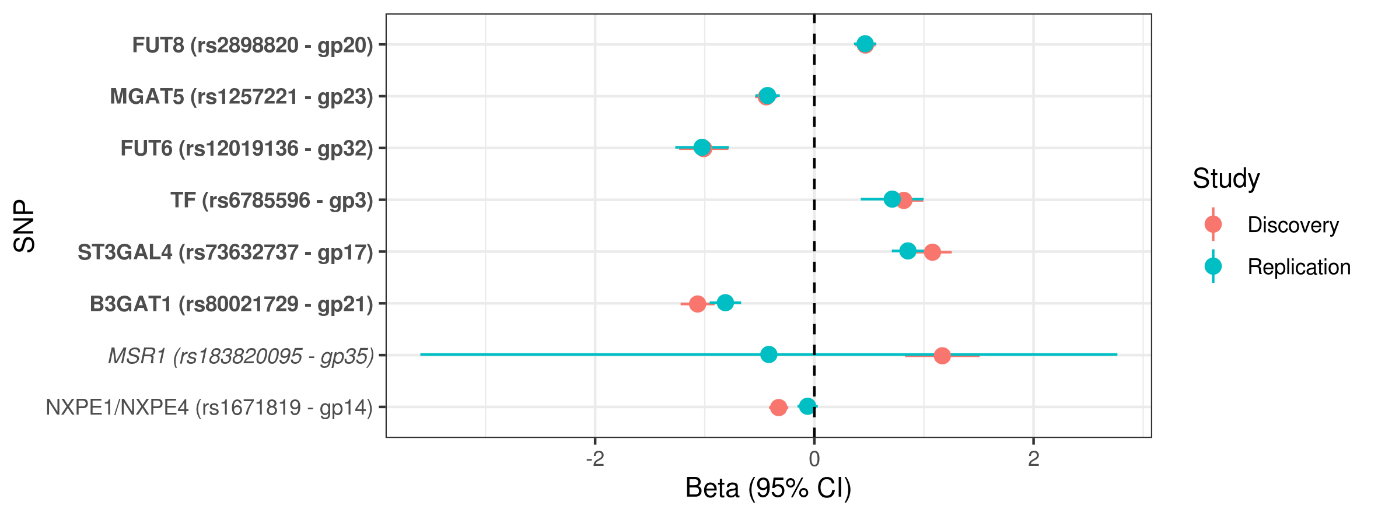

**Supplementary Figure 6. Replication of discovery GWAS.** For each locus associated with transferrin N-glycome in CROATIA-Korcula cohort, effect size and standard error of top SNP are reported in red, when estimated in CROATIA-Korcula cohort (Discovery), and in blue, when estimated in VIKING cohort (Replication). Gene names have been marked by different fonts based on overlap between confidence intervals (CIs) of effect estimates. **In** **bold:** nominal replication (p < 0.05). *In* *italics:* CIs overlap and cover zero, and replication estimate is closer to zero than discovery. In roman: CIs do not overlap and replication estimate covers zero. Proxy SNP rs554715390 was used for replicating SNP rs183820095 (D’=1, r^2^=1).
